# *In vivo* dissection of human *NRXN1* isoforms reveals gain-of-function pathogenicity of schizophrenia-associated 3’ deletions

**DOI:** 10.1101/2025.08.29.672776

**Authors:** Dustin Haskell, Michael P. Hart

**Author notes:** corresponding author Michael P. Hart –.

## Abstract

Heterozygous deletions in *NRXN1*, encoding the presynaptic adhesion molecule Neurexin 1, are among the most frequently identified rare variants in schizophrenia and other neuropsychiatric disorders. Patient sequencing has revealed that 3’ deletions within *NRXN1* generate novel isoforms not produced from the intact locus, yet whether these isoforms are functional, passively non-functional, or actively pathogenic *in vivo* is still not clear. To address this, we expressed eight human *NRXN1* isoforms in *C. elegans* neurons including four control isoforms and four 3’ deletion variant isoforms identified in schizophrenia patient cell lines, and characterized their effects on protein localization and two independent behaviors: a food deprivation response and social feeding behaviors. Most human isoforms showed expression and localization within the nerve ring and neurons similar to the *C. elegans* ortholog, NRX-1; however, several isoforms, particularly among the 3’ deletion variants, displayed aberrant accumulation in neuronal cell bodies or as puncta in neuropil. Functionally, isoforms fell into one of three categories: no effect on *nrx-1* loss-of-function behavioral phenotypes, partial rescue, or gain of function, with multiple isoforms showing differences between the behaviors. Strikingly, two 3’ deletion isoforms produced gain-of-function behavioral phenotypes more severe than the *nrx-1* null mutant, demonstrating that these patient-derived variants can be actively pathogenic. These results establish *C. elegans* as a tractable *in vivo* platform for dissecting the isoform-specific functional consequences of *NRXN1* variants and suggest that strategies for *NRXN1*-associated neuropsychiatric diseases must account for both loss-of-function and gain-of-function isoform mechanisms.

## Introduction

The evolution of complex behavioral traits came about through selective pressures across distinct and diverse populations through adaptation in neurodevelopmental and neuronal functional traits^1^. Strongly damaging mutations and variants in genes essential for neurodevelopment are generally selected against, while in contrast, less damaging or partially penetrant variants can result in a range of phenotypic severity^1,2^. Selective pressures on genes with less severe variants can modulate non-essential behavioral traits and drive evolutionary changes in behaviors by modifying development and function of the brain^3^. As a consequence of these adaptations, some behavioral changes that arise in these unique evolutionary trajectories can include those associated with neuropsychiatric disorders^4^. However, the complex genetic and biochemical landscape of human neurodevelopment, which includes incredibly diverse neuronal subtypes each with unique developmental trajectories, has obscured our understanding of the molecular and neuronal underpinnings of behaviors observed in neuropsychiatric disorders like schizophrenia.

To understand the network of neuronal genes driving neurodevelopment and neuropsychiatric disorders, case control and large gene association studies have been carried out to identify genes associated with risk for disorders diagnosed by behavioral changes, including schizophrenia ^5–8^. Interestingly, there is significant overlap of select genes (and similar perturbations) implicated across multiple conditions and disorders defined by shared and distinct behavioral changes^9–12^. These decisive nature of these risk factor genes raise many questions, for example, What are the roles of these genes at the neuron and circuit level that make them essential for development of many behaviors? Do different variants within a single gene lead to specific neurodevelopmental trajectories or functional changes that result in behavioral changes associated with different disorders? Overall, to understand how specific changes in a singular gene or across gene cohorts associated with a disorder can lead to alterations in disparate behaviors will require a mechanistic understanding of how variants alter function of genes, ideally linked to behavioral changes.

To begin to answer some of these questions, one tractable approach is to focus on a subset of associated genes and variants^5,11^. Synaptic adhesion molecules are essential for synaptogenesis, organization, and function by establishing synaptic connectivity and polarity through protein-protein interactions, and thereby unrepining critical behavioral functions^13–15^. Among the best studied synaptic adhesion molecules are neurexins (NRXN). Mammals have three neurexin genes, *NRXN1-3*, from which two separate promoters generate a long α-isoform and a shorter β-isoform^16–18^. Additionally, *NRXN1* also produces a very short γ-isoform from an overlapping and internal promoter^18^. In contrast to mammals, *Drosophila* and *C. elegans* encode a single neurexin orthologue^19,20^, with the *C. elegans* gene *nrx-1* encoding a long α-isoform and short γ-isoform similar to mammalian *NRXN1*^21^.

Like many transmembrane synaptic adhesion proteins, neurexins are comprised of a large extracellular domain containing multiple domains for protein-protein interactions as well as a short intracellular domain for synaptic scaffolding and signaling^13,22^. Historically, neurexins act primarily with a cognate binding partner neuroligins to establish pre-synaptic and post-synaptic polarity^23,24^. Early work was consistent with both proteins being essential for Ca^2+^-dependent neurotransmission at both excitatory and inhibitory synapses^25–27^. Later work clarified that neurexins’ interactions and roles at the synapse are far more complex, with the protein showing tissue, cell type, and even synapse-type specific mechanisms. Many additional binding partners have been subsequently identified (including neurexophilin, dystroglycan, LRRTM proteins, and cerebellin) which greatly expand the possible mechanisms through which neurexins act^17,28^. From a structural perspective neurexin is relatively simple, with the α-isoform having six LNS domains periodically interspersed with EGF-like domains. However, additional complexity arises from a number of commonly utilized alternative splice sites (five in the α-isoform and 2 in the β-isoform of *NRXN1*), potentially generating over 1,000 possible splice variants^29^. However, RNAseq data suggests that only a relatively small subset of the total isoform permutations (∼200 alpha isoforms) are consistently expressed in any given tissue or cell type^30–32^. Detailed mechanistic study of these splice sites uncovered several roles for alternative splicing of neurexins in modulation of binding affinity to specific ligands and partners like neuroligin, as well as controlling the ratio of neurexin isoforms in a given synapse^33^.

Initial characterizations of the three mammalian neurexins suggested similarity in functions and redundancy, however, tissue heterogeneity, developmental differences, and isoform/splice variants have demonstrated a complexity in the functions of each neurexin gene and even isoforms^13^. In contrast, the single neurexin orthologue in *C. elegans* and flies is well conserved and provides a model to study the mechanisms and effects of neurexin. Given that neurexin has many important roles in synaptic development, function, and maintenance, it is not surprising that genetic variation across neurexin genes is associated across many neuropsychiatric disorders and conditions. For example, perturbations ranging from point mutations to large exonic deletions in *NRXN1* have been associated with autism, Tourette’s, bipolar disorder, suicide risk, and schizophrenia^34–39^. Recent work has shown copy-number variants of neurexin contribute significantly to the risk of schizophrenia^37^. Furthermore, heterozygous exonic deletions are consistently found in patients with schizophrenia, some of which have been implicated in the impairment in neurotransmitter release and neuronal development, as well as an overall reduction in neuronal activity^40^. However, how individual and often disparate variants in the *NRXN1* gene impact its function to ultimately result in behavioral differences is not well understood, in part due to the complexity of neurexin isoforms, which can be differentially impacted by variants across the gene length. Even large deletions in *NRXN1*, which are traditionally thought to introduce loss of function of specific isoforms, were found to result in generation of novel isoforms not generated from the intact locus.

Here we used *C. elegans* to explore the impact of *NRXN1* isoforms derived from a patient harboring a 3’ deletion on NRXN1 expression and localization, as well as neuronal function and seveal stereotyped behaviors. First, we found that human *NRXN1* isoforms can be expressed in *C. elegans* neurons, with some isoforms showing expression and localization similar to expression of the NRX-1 alpha isoform, while some showed distinct punctate or accumulation in neuron cell bodies. Next, we found that some control *NRXN1* isoforms were able to rescue behavioral phenotypes observed with loss of *nrx-1*, including food deprivation response and social feeding behavioral phenotypes, while other control isoforms having no impact on behavior. Meanwhile the novel *NRXN1* isoforms generated from the 3’ deletion differentially impacted behaviors, some partially rescued behavioral phenotypes observed with loss of *nrx-1*, some had no impact, and others showed gain of function phenotypes, indicating they may impart some level of pathogenicity. While most *NRXN1* isoforms show similar impacts across both behaviors, several showed distinct differences between the behaviors, suggesting behavior-specific functional roles for *NRXN1* isoforms, with or without disease-derived variations.

## Methods

### Cloning

*NRXN1* isoform cDNAs were acquired in lentiviral plasmids from the lab of Kristen Brennand (Yale)^41^. To generate plasmids for expression of human *NRXN1* in *C. elegans, NRXN1* cDNAs were cloned into pMPH34 (p*ric-19::sfGFP::nrx-1(α)*)^42^ by Epoch Life Sciences, replacing the existing *C. elegans nrx-1*α cDNA. This plasmid drives expression in all neurons using the *ric-19* promoter and a neutral *unc-54* 3’ UTR. To remove the N-terminal signal sequence of human *NRXN1*, RF cloning was utilized (all plasmids are listed in **Supplemental Table 1**). After cloning all plasmid sequences were confirmed via Sanger Sequencing.

### *C. elegans* Strain Construction and Maintenance

Animals were maintained under normal growth conditions (∼20° C) on normal NGM media^43^. Plates were seeded with *E. coli* OP50 to provide nutrition. *NRXN1* isoform transgenic animals were generated by injecting the human *NRXN1 C. elegans* expression plasmids for each *NRXN1* isoform. Standard procedure was used for microinjections. *NRXN1* isoforms were injected at a concentration of 40 ng/μL along with 20 ng/μL of a co-injection marker (either p*lin-44::gfp* or p*unc-122::mScarlet*). Injections were done into the null allele *nrx-1(wy778)*. Once stable extrachromosomal lines were established and characterized for transmission and variability, transgenes were crossed into the *npr-*1*(ad609)* background for the social feeding assays. All strains used are listed in **Supplemental Table 1**.

### Food deprivation behavioral response and activity assays

Animal locomotion and activity levels were analyzed using the WorMotel device as previously described^42,44^. Briefly, each of the 48 wells in the WorMotel chip were filled with standard NGM and allowed to cool. For food deprivation conditions, young adult animals were picked off a plate into a well of M9 media to remove any residual bacteria from their cuticle. Each animal was then carefully pipetted in 1μL of M9 and placed in a WorMotel well. Once all the animals had been pipetted the liquid was allowed to fully dry before starting the behavioral assay. The WorMotel was placed in a 100 mm petri dish humidified with a wet kimwipe to prevent desiccation.

The WormWatcher imaging platform was utilized to capture images as previously described^44^, with images being captured every 10 seconds for 8 hours. Previously published custom MATLAB code was used to process and analyze the images. The total pixels displaced over a 1-hour period was calculated for each well. Images were manually inspected to remove any wells with escaped, sickly, or stuck animals which were excluded from final analysis. Each was run (3-5) individual replicates, which were combined.

### Social feeding behavioral assay

To assay social feeding behavior, 6-well culture plates were filled with NGM media (6mLs) and seeded with 75 μL of *E. coli* OP50 and allowed to dry overnight. To assess levels of social feeding aggregation, 50 L4 animals were picked onto the seeded wells. To prevent condensation during imaging, a small amount of Tween20 was used to coat the surface of the 6-well lid. The plates are imaged using the WormWatcher automated imaging platform (Tau Scientific) for a total of 20 hours with a cluster of 10 images (over a 1-minute timecourse) being taken every hour. Quantification of social feeding aggregation was performed as previously described^45^. Briefly, an animal is considered aggregating if it is in contact with two or more other animals.

### Microscopy

Young adult animals were anesthetized in ∼4 μL of 100 μM sodium azide before being immobilized on a standard 5% agarose pad and coverslip. A Leica inverted TCS SP8 laser-scanning confocal microscope with 60x objective lens was used to capture the head regions of each animal to analyze the expression and localization of each *NRXN1* isoform within the nerve ring surrounding the pharynx. Z-stacks were set at 0.5 microns, and Z-imaging captured the full Z-plane of tissue. Consistent laser power and gain settings were used in order to standardize analysis of the expression patterns and levels. Micrographs were generated by generating a SumStack of the available Z-stacks to produce a compressed expression pattern in the nerve ring.

### Statistics

The number for each experiment was based on previous studies and effect size, with each experiment performed with at least 3 independent replicates and each trial performed with matched controls^42,45^. All data were analyzed and plotted in GraphPad Prism 10 and statistical significance was determined using one-way ANOVA with Tukey’s post-hoc test. For comparisons of two data sets, a two-tailed unpaired t-test was used to compare significance. Error bars on figures represent standard error of the mean (SEM) and p-values are shown in each figure to indicate significance (P<0.05).

## Results

### Human *NRXN1* isoform expression and localization in *C. elegans* neurons

To analyze potential differences across different *NRXN1* isoforms, we first compared each schizophrenia-derived isoform to the longest α-isoform of human *NRXN1*. Mapping the 3’ deletion onto the genomic locus and protein domain architecture demonstrates that all 4 of the 3’ deletion isoforms excise exons 20 and 22 at the C-terminus of the protein **(Figure 1A**). Furthermore, the deletion is localized to the extracellular region of the protein potentially affecting both localization within the membrane and flexibility of the extracellular domains. Removal of exons 20/21 in the 3’ deletion isoforms significantly reduce the number of unstructured residues present between the transmembrane domain and the globular domains (**Figure 1B**). This structural change has potential functional consequences: the linker region between the transmembrane domain and the LNS6/EGF3 globular domains contributes to the conformational flexibility that allows the neurexin ectodomain to adopt the correct orientation in the synaptic cleft for binding to postsynaptic ligands such as neuroligins, LRRTMs, and cerebellins. Reduction in length of this unstructured region may therefore impair ectodomain orientation or intracellular trafficking, and potentially disrupt expression, localization, and functional interactions.

**Figure 1.**
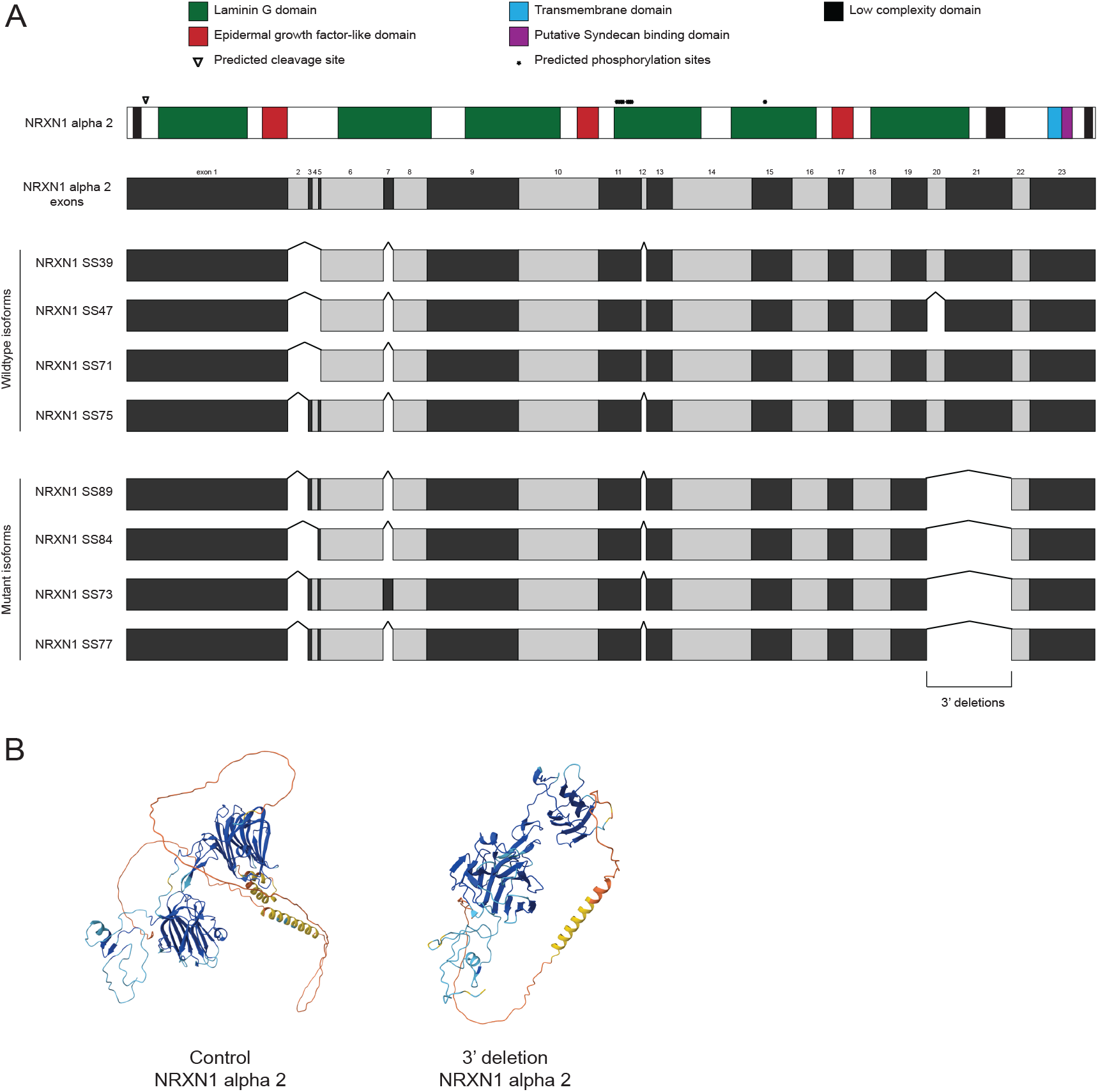
Schematic of NRXN1 isoform variants. **(A)** Eight NRXN1 isoforms aligned against the *NRXN1* α2 (representing the longest alpha isoform) isoform protein domain architecture. Normal splice variations are represented by interruptions within the CDS, while 3’ NRXN1 deletions present in the schizophrenia-derived isoforms are labeled. (B) Alphafold predictions of exons 14-23 of control NRXN1 α2 (left) and 3’ deletion SS84 (right).

To better understand the functional impact of the 3’ deletion on each isoform, we expressed the human *NRXN1* isoforms in all neurons of *C. elegans*. Aligning the longest α-isoform of human NRXN1 and the 4 control NRXN1 isoforms to the longest *C. elegans* NRX-1 isoform confirms conservation of the protein domains and architecture, with a few short human and *C. elegans* specific protein sequences, including a number of exons differentially included in the human isoforms (**Supplemental Figure 1**). To make ‘humanized’ *NRXN1 C. elegans*, we first generated expression plasmids where the cDNA for each human NRXN1 isoform was inserted to replace the *nrx-1* cDNA in a previously described *nrx-1*α expression plasmid, which includes the *ric-19* promoter to drive expression in all neurons, an artificial intron, signal sequence, and a superfolder *gfp* tag with a linker to the respective neurexin cDNA^42,45^. We then injected each *NRXN1* isoform plasmid into animals with a large deletion in the single ortholog of neurexin (*nrx-1*) that removes the majority of the alpha and all of the gamma isoform coding regions, and acts as a functional null (*nrx-1(wy778))*. We observed expression of all of the superfolder *gfp* tagged human *NRXN1* isoforms, which had expression patterns (and levels) comparable to the same transgenic expression of *nrx-1*^42,45^, with localization to neurons and the nerve ring (**Figure 2A**). Expression of endogenous NRX-1α with the same tag shows a diffuse punctate pattern of expression in the nerve ring neuropil and extends ventrally and into the nerve cord^42^. Typically, there is only dim expression in the cell bodies of the head neurons or within projections extended towards the nose^42^, which was observed for SS71 and SS75 (**Figure 2A**). Several of the human isoforms (particularly SS47 and SS73, representing the control and 3’ deletion versions of a single isoform, respectively) showed some ectopic expression in the cell bodies of head neurons. The SS73 variant showed localization primarily to the cell bodies of many neurons, with fainter and diffuse expression in the axons in the nerve ring (**Figure 2B**). Lastly, the SS77 and SS89 isoforms showed a more punctate expression in both the cell bodies and/or axonal projections of the nerve ring that was not observed in the other isoforms (**Figure 2B**).

**Figure 2.**
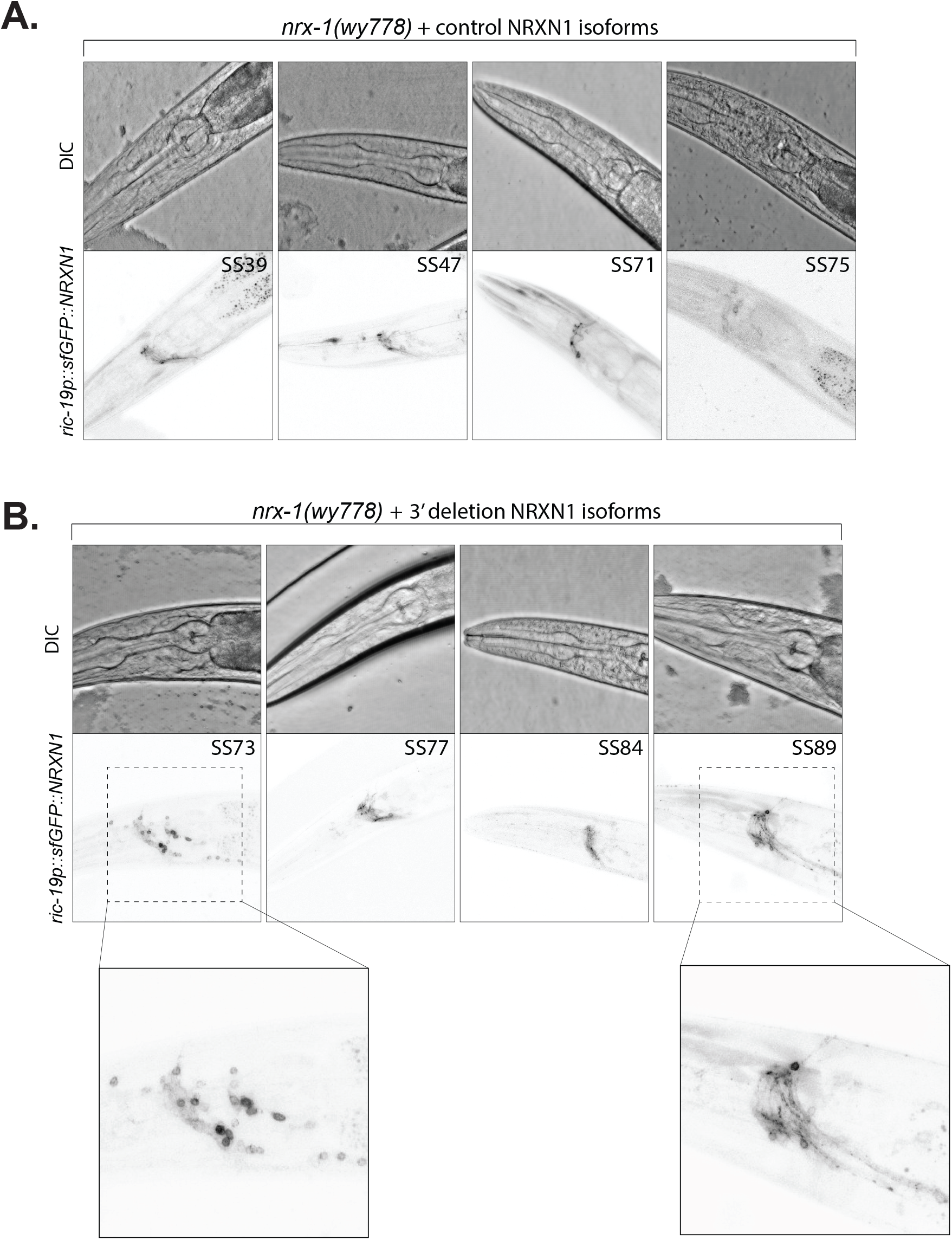
Expression and localization of NRXN1 isoforms in *C. elegans* neurons. **(A)** *ric-19p*:*:sfgfp::NRXN1* drove expression of control NRXN1α isoforms (top panel), with the majority of the expression being localized to nerve ring surrounding the pharynx. The lower panel is DIC for anatomical reference. **(B)** *ric-19p*:*:sfgfp::NRXN1* drove expression of 3’ deletion isoforms of NRXN1α (top panel), with the majority of expression localized to nerve ring surrounding the pharynx. Insets show some expression within cell bodies of head neurons, and some abnormal punctate expression of human NRXN1 isoforms.

**Supplemental Figure 1.**
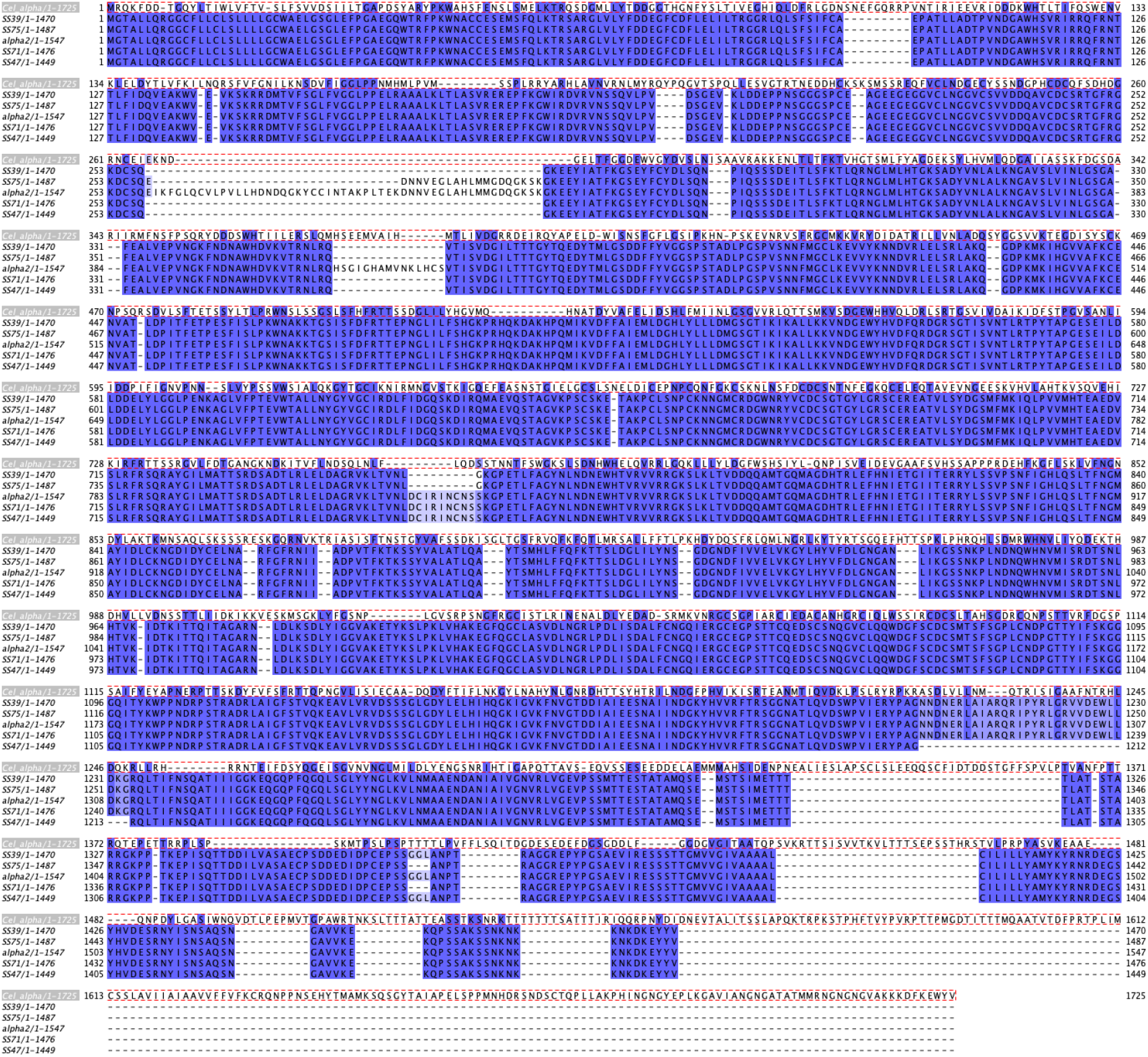
Multiple sequence alignment of NRXN1 alpha 2 isoform, wildtype NRXN1 variants and *C. elegans* NRX-1 alpha isoform. CLUSTAL omega alignment of neurexin proteins with percent identity of conserved residues highlighted in blue.

### *NRXN1* isoforms differentially impact activity levels of food-deprived animals

We next tested the functional impact of each NRXN1 isoform on neuronal and circuit function, using behavioral phenotypes we previously described for loss of function of *nrx-1*^42,45^. This previous work showed that *C. elegans* have a robust response to food deprivation, characterized by a significant and sustained increase in their activity levels compared to animals with food^42^. Several genes show behavioral phenotypes in this behavioral assay, including *nrx-1* mutants, which have a significantly reduced activity upon food deprivation. Isoform-specific alleles and transgenic experiments showed that the γ-isoform and the α-isoform of *nrx-1* play distinct temporal roles in the initiation and maintenance of hyperactivity upon food deprivation^42^.

To test whether human *NRXN1* isoforms could rescue the *nrx-1(wy778)* behavioral phenotype, we quantified activity levels of animals without food using the WorMotel^42,46^. In comparison to control animals (N2), the *nrx-1(wy778)* animals showed a significant decrease in activity (**Figure 3A**). Expression of human *NRXN1* isoforms differentially impacted activity levels *of nrx-1(wy778)* animals. Two control isoforms (SS71 and SS75) increased activity and displayed partial rescue the *nrx-1(wy778)* phenotype, suggesting that they can functionally compensate for the loss of NRX-1 (**Figure 3A, left panel**). The other two control isoforms (SS39 and SS47) had no impact on activity levels of *nrx-1(wy778)* animals (**Figure 3A**). In comparison, none of the 3’ deletion *NRXN1* isoforms increase activity levels or rescue the *nrx-1(wy778)* phenotype. Interestingly, two of the 3’ deletion isoform variants (SS77 and SS89) showed a gain of function phenotype, where activity levels were significantly reduced compared to the *nrx-1* null animals (**Figure 3A, right panel**). Overall, human NRXN1 isoforms displayed different functional impacts on the food deprivation behavioral phenotype: partial rescue (suggesting evolutionary conservation of function)(**Figure 3B**), no impact (where the activity levels were similar to *nrx-1(wy778)*)(**Figure 3C**), and gain of function (where activity levels were reduced further than loss of *nrx-1* alone (**Figure 3D**).

**Figure 3.**
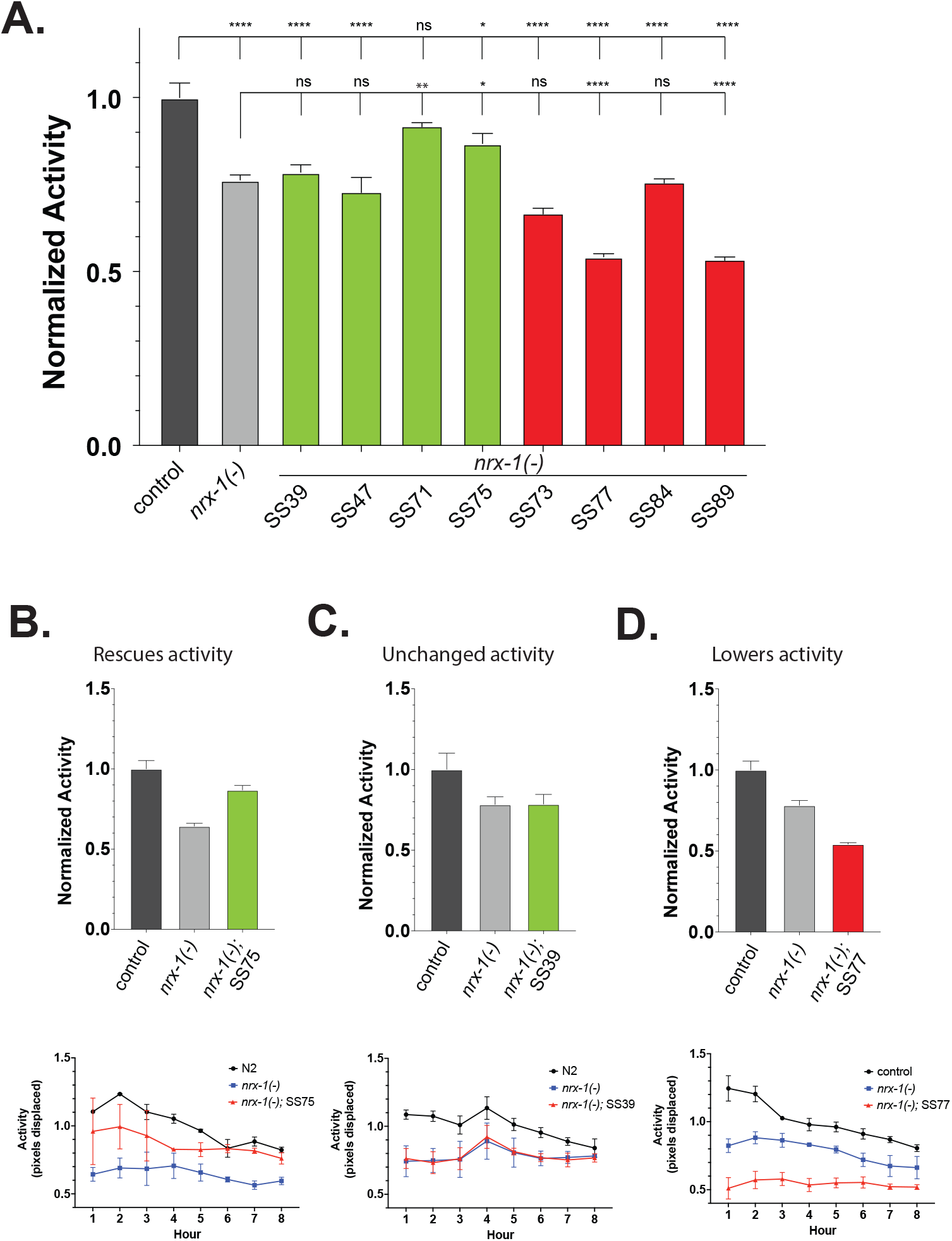
Comparison of NRXN1 isoforms on food-deprived activity levels. **(A)** Normalized activity values for animals without food. All *nrx-1(wy778)* and *NRXN1* animals were normalized within replicates to allow for pooling of replicates across days and equipment setups. A one-way ANOVA, with Tukey’s post-hoc test was utilized to determine statistical significance. (B) SS75 is shown as an example of an isoform variant that is able to increase activity of the *nrx-1(wy778)* animals. Top panel shows normalized average across the full 8-hour time course. Bottom panel shows the 8-hour time series broken down by hour. **(C)** SS39 is shown as an example of an isoform variant whose activity is unchanged compared to the *nrx-1(wy778)*. Top panel shows normalized average across the full 8-hour time course. Bottom panel shows the 8-hour time series broken down by hour. **(D)** SS77 is shown as an example of an isoform variant whose activity is significantly lower compared to the *nrx-1(wy778)*, thus representing a gain of function phenotype. Top panel shows normalized average across the full 8-hour time course. Bottom panel shows the 8-hour time series broken down by hour.

The partial rescue of *nrx-1* behavioral deficits by SS71 and SS75 isoforms is itself a significant finding: it demonstrates functional conservation of core neurexin signaling properties across approximately 700 million years of evolution, validating the use of *C. elegans* as an *in vivo* model for human NRXN1 variant biology. Equally notable is that two intact human control isoforms (SS39 and SS47) showed no rescue despite being full-length, wild-type sequences, demonstrating that isoform functional divergence exists among control isoforms independently of any deletion. This finding suggests that the specific combination of splice site variants defining each isoform determines its capacity to engage the neuronal circuits governing these behaviors, consistent with neurexin isoforms carrying a molecular code for circuit-specific synapse properties.

### NRXN1 isoforms differentially impact social feeding behaviors

To further characterize the *NRXN1* isoforms, we also analyzed their functional impact on social feeding behavior, specifically the aggregation of *C. elegans* in groups while feeding^45,47^. To perform this assay, all of the *NRXN1* isoform transgenic *nrx-1(wy778)* strains were crossed into the *npr-1(ad609)* background, which serves as a genetic manipulation to induce aggregation behavior observed in wild isolate *C. elegans* in the lab N2 Bristol strain (which shows solitary feeding behavior)^47^. We previously showed that both a complete null or the specific loss of the α isoform of *nrx-1* are sufficient to strongly reduces social feeding behavior. These defects can be then be rescued by expression of NRX-1 α-isoform in all neurons or subsets of neurons^45^, and thus represents a sensitive assay to test *NRXN1* functionality. First, we tested the impact of *NRXN1* isoforms on social feeding behavior in solitary control background (*nrx-1(wy778))* and observed no significant aggregation in control animals or animals expressing the 8 human *NRXN1* isoforms (**Supplemental Figure 2**). However, we found that expression of the *NRXN1* isoforms in the social control *npr-1(ad609)* background induced a range of impacts on social feeding behavior (**Figure 4A**). We confirmed that *npr-1(ad609)* social feeding controls have high levels of aggregation (averaging around 40/50 animals), whereas the *nrx-1(wy778); npr1-(ad609)* control has a significantly lower level of aggregating animals **(Figure 4A&B**). The *NRXN1* control isoforms had variable impact, where 3 control isoforms did not impact levels of aggregation compared to *nrx-1(wy778); npr1-(ad609)* animals, but SS75 increased aggregation to the point of partially rescuing the *nrx-1* phenotype (**Figure 4A, left panel**). Interestingly, although 3 of the 3’ deletion NRXN1 isoforms showed increased aggregation compared to *nrx1(wy778); npr1-(ad609)*, SS73 represents a significant rescue of the aggregation phenotype, with levels indistinguishable from *npr-1(ad609)* levels (**Figure 4A, right panel**). Additionally, we found that SS89 displayed a strong gain of function phenotype, with aggregation significantly lower than the *nrx-1(wy778); npr1-(ad609)* control (**Figure 4A, right panel**), similar to its impact in the food deprivation response behavior.

**Figure 4.**
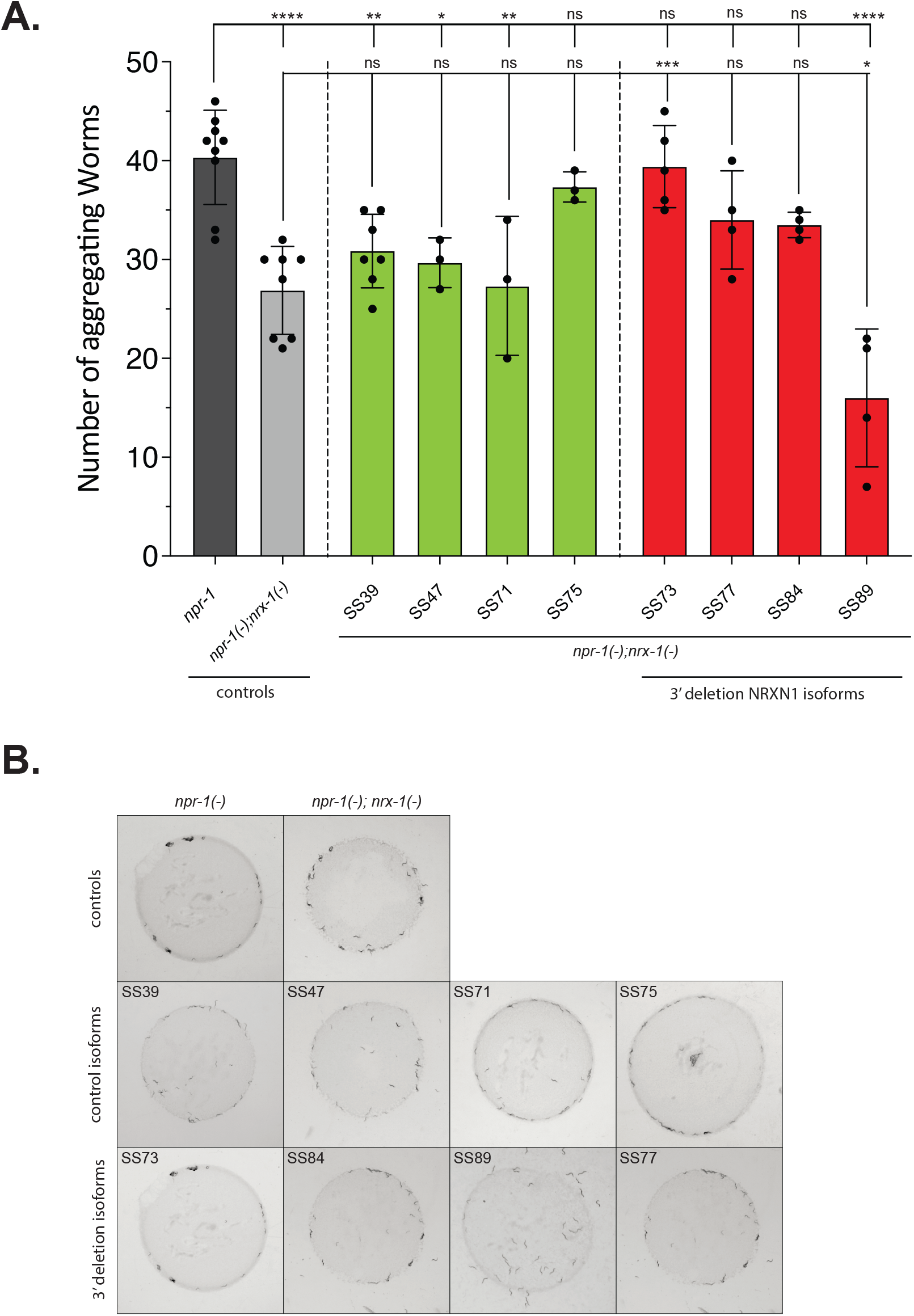
NRXN1 isoforms differentially impact social feeding behavior. **(A)** In comparison to *npr-1(ad609)* controls with high aggregation, *npr-1(ad609);nrx-1(wy778)* controls have significantly reduced levels of aggregation. Control NRXN1 isoforms are shown in the left panel, while 3’ deletion isoform variants are shown on the right panel. Bar graphs represent the mean values across biological replicates (each replicate indicated by a dot). Error bars show stand error of mean. One-way ANOVA with Tukey’s post-hoc test was used for statistical comparisons, with p-values indicated in standard format (p=0.05(*), p=0.01(**), p=0.001(***), p= <0.001 (****)). **(B)** Representative images of aggregation behavior for each genotype.

Importantly, several isoforms showed quantitatively different or discordant effects between the two behavioral assays. This behavior-specific variation is consistent with *nrx-1* isoforms having circuit-specific roles in *C. elegans*: the food deprivation response is mediated in part through the octopamine-producing RIC neuron and monoamine signaling, while social feeding aggregation depends on *nrx-1* function in two pairs of glutamate sensory neurons^42,45^. The differential sensitivity of these circuits to specific isoforms suggests that individual NRXN1 splice variants may have circuit-selective functions even in *C. elegans*, a finding that will be important to consider when modeling the behavioral consequences of specific isoform repertoire changes in patients. Notably, the isoforms displaying aberrant cell body-enriched localization (SS73 and SS89) were precisely those that showed gain-of-function behavioral phenotypes, while isoforms with nerve ring-predominant expression comparable to NRX-1 (SS71, SS75) were those capable of partial behavioral rescue. This correspondence between localization pattern and functional outcome suggests that proper synaptic targeting is a prerequisite for NRXN1 function in these behavioral circuits.

**Supplemental Figure 2.**
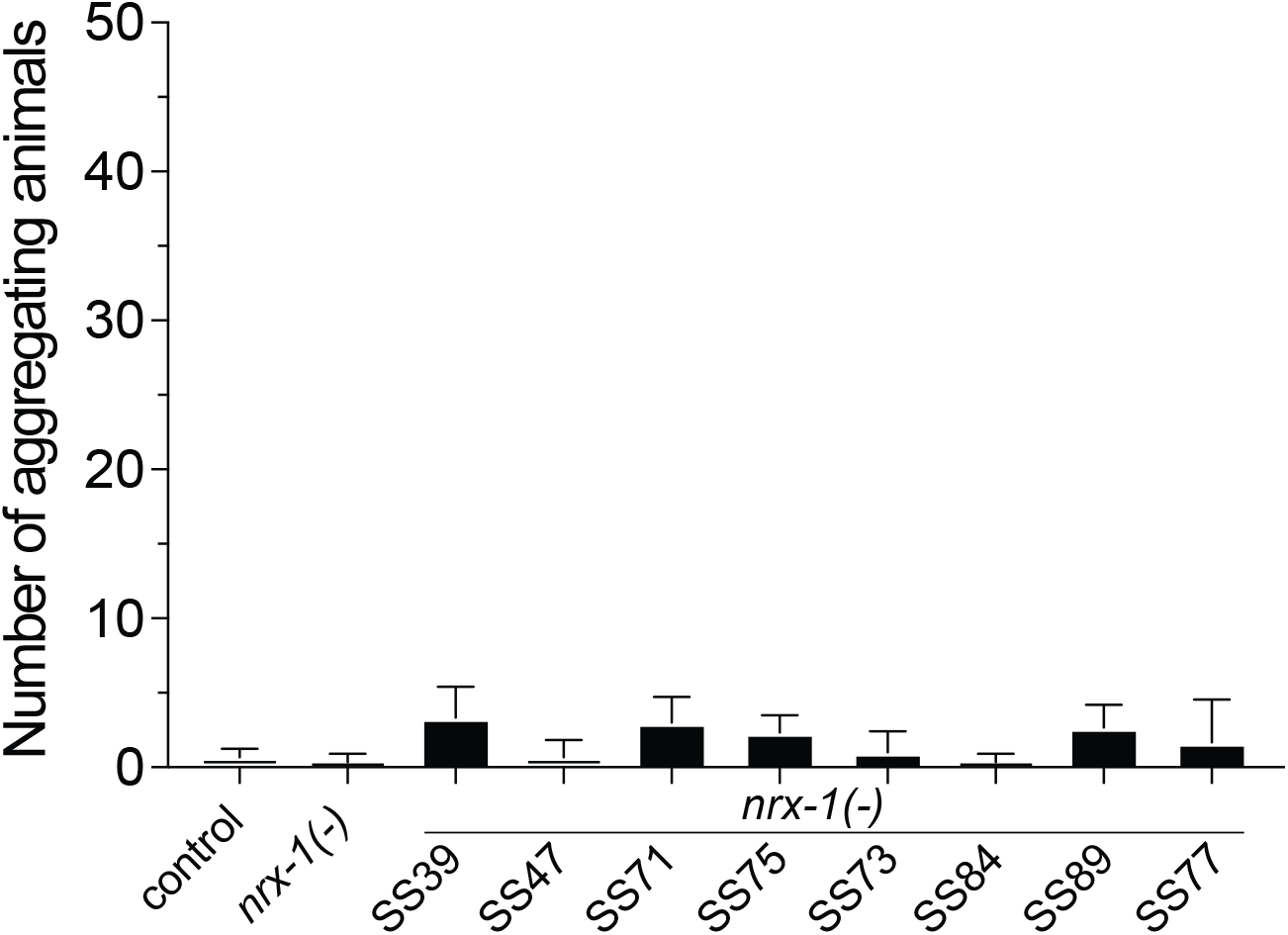
Expression of NRXN1 isoforms does not induce aggregation behavior. In addition to control (N2 Bristol) and *nrx-1(wy778)*, 8 NRXN1 isoforms (4 control isoforms and 4 3’ deletion isoforms) were tested for aggregation behavior in the solitary feeding N2 Bristol strain background with *nrx-1(wy778)*(as opposed to the social feeding and aggregating *npr-1(ad609)* background).

## Discussion

Synaptic adhesion proteins including NRXN1 are widely regarded as essential drivers of synaptic assembly and organization as well as maintenance of overall synaptic function. While small perturbations or partial loss of the neurexin family in mammals is generally tolerated^27^, these changes can and are implicated in a number of developmental, neuronal, and behavioral changes representing a rage of serverity^13,40,41^. In parallel, with the creation of large genotyping and genome sequencing projects, it has emerged that heterozygous insertions/deletions, point mutations, and copy number variants in *NRXN1* contribute to its association with various human conditions, indicating that these variants impact protein function at the mechanistic level^13,17,35–39^. Consolidating these additional implication of variant-specific molecular effects with the immensely complex Neurexin splicing network provides a highly intricate picture of neurexin function.

In this work, we characterized the impact of expressing eight NRXN1 isoforms (4 control and 3’ deletion variants each) isolated from schizophrenia-derived hiPSCs^41^. These isoforms were identified as among the most highly expressed NRXN1α isoforms in patient-derived neurons, and span multiple alternative splice site combinations. This cohort includes isoforms whose levels are substantially reduced from the wild-type allele and novel isoforms generated *de novo* from the deletion allele^41^. Leveraging the benefits of *C. elegans* has allowed for the detection of both molecular and behavioral differences between control and 3’ deletion variants of individual isoforms, ultimately showing that even small isoform differences can significantly alter behavioral outputs. We observed differential expression patterns across NRXN1 isoforms, providing insight into potential mechanisms by which the 3’ deletion may affect protein function. Proper presynaptic localization of neurexins are essential for their roles in engaging postsynaptic ligands and organizing the synapse, meaning that protein sequestered in the neuronal cell body or neuropil puncta cannot fulfill these functions. The 3’ deletion shared across all four isoforms removes exons 20 and 21, which our AlphaFold modeling predicts significantly reduces the unstructured linker between the transmembrane domain and the extracellular globular domains. This structural constraint may impair correct ectodomain folding or trafficking, causing the protein to accumulate in the cell body or puncta in the neuropil rather than traffic efficiently to synapses. Isoforms with primarily nerve ring localization at dimmer levels (SS71, SS75) were those capable of behavioral rescue, with more punctate expression inducing gain-of-function behavioral phenotypes (SS77, SS89)(**Table 1**). This correlation between localization pattern and functional outcome supports the hypothesis that synaptic targeting is a prerequisite for NRXN1 function in these behavioral contexts. Specific ectopic localization may itself contribute to behavioral pathology, as mis-localized presynaptic proteins have been shown to disrupt neuronal protein homeostasis.

**Table 1.** Summary of localization and behavioral phenotypes of human NRXN1 isoforms.

| **NRXN1 isoform** | **localization** | **activity off food** | **social feeding** |
| --- | --- | --- | --- |
| SS39 | normal | / | / |
| SS47 | normal, soma | / | / |
| SS71 | restricted | increase | / |
| SS75 | reduced | increase | trend up |
| SS73 3' del | mostly cell soma | / | increase |
| SS77 3' del | punctate | decrease | trend up |
| SS84 3' del | normal | / | trend up |
| SS89 3' del | more punctate | decrease | decrease |

Particularly striking was the gain-of-function observed for two of the 3’ deletion isoforms, SS77 and SS89, in the food deprivation assay, with SS89 also showing a strong gain-of-function in social feeding behavior (**Table 1**). These isoforms did not simply fail to function, they produced activity levels or aggregation phenotypes more extreme than the null mutant alone. This gain-of-function could reflect several non-mutually exclusive mechanisms: (1) a dominant-negative effect, in which the truncated NRXN1 protein competes with endogenous NRX-1 for binding to postsynaptic partners, sequestering ligands in a non-functional complex; (2) a neomorphic function, in which the structural change introduced by removal of exons 20/21 creates a novel interface for protein-protein interactions and thus engages aberrant signaling pathways; or (3) toxic effects arising from the accumulation of misfolded or mislocalized protein in the cell body. Our results complement findings from recent studies that demonstrated using patient-derived hiPSC neurons that *NRXN1* heterozygous deletions operate through both loss-of-function and gain-of-function mechanisms in a cell-type-specific manner, with gain-of-function effects particularly prominent in GABAergic neurons^40,41^. Our findings in *C. elegans* are consistent with and extend this framework, providing *in vivo* behavioral evidence that truncated NRXN1 isoforms generated by 3’ deletions can have pathogenic functions beyond simple loss of wild-type activity. From a genotype perspective, this distinction is critical: a loss-of-function model would predict that restoring wild-type NRXN1 expression is sufficient for therapy, whereas a gain-of-function model requires additionally eliminating the pathogenic isoform, highlighting a need for genotype-specific interventions for NRXN1 variant carriers. The requirements may differ between patients depending on the specific deletion, the isoform repertoire expressed in affected cell types, and the balance of isoforms at the patient’s locus, underscoring the importance of isoform-level characterization in both research and clinical contexts.

The ability of human NRXN1 isoforms to partially rescue *C. elegans* behavioral phenotypes is remarkable given the approximately 700 million years of evolutionary divergence between humans and nematodes. This cross-species functional conservation underscores the deep evolutionary context of neurexin’s core synaptic roles and validates *C. elegans* as an *in vivo* model for studying the functional consequences of human NRXN1 variants. However, the partial nature of rescue even for the most active isoforms (SS71 and SS75) is expected given the species-specific differences in downstream signaling partners, neuronal circuit architecture, and the context of isoform repertoires that may limit full complementation in a heterologous system. An important observation from this work is that the effects of NRXN1 isoforms were not entirely consistent across the two behavioral assays: some isoforms showed behavioral effects of differing magnitude between the food deprivation and social feeding paradigms. This behavior-specific variation is consistent with the growing body of evidence that neurexin isoforms can have neuron-, circuit-, and context-specific roles. In *C. elegans*, the alpha and gamma isoforms of *nrx-1* show distinct temporal contributions to the food deprivation response and in chemosensory responses, and neurexin’s role in social feeding is mediated through specific neuronal subtypes and downstream monoamine signaling pathways^42,45,48^. The differential sensitivity of these two behavioral assays to specific NRXN1 isoforms may reflect differences in which neuronal circuits or synaptic partners are most critical for each behavior, and future work identifying the neurons and molecular pathways most sensitive to specific isoform variants could provide mechanistic insight.

Several important limitations of this study should be acknowledged. First, the NRXN1 isoforms were expressed as extrachromosomal arrays, which introduces copy number variability and mosaic expression across animals. While we controlled for this by characterizing and selecting transgenes from multiple independent lines, using matched controls, and replicating all experiments, future work using single-copy chromosomally integrated transgenes (e.g., via MosSCI or CRISPR knock-in) would provide more precise and quantitative comparisons between isoforms. Second, all NRXN1 isoforms were expressed under a pan-neuronal promoter; endogenous human NRXN1 shows highly regulated, cell-type-specific expression that may be important for its specific functions and interactions. Third, the behavioral assays used here capture circuit-level outputs and do not directly measure synaptic structure or function; complementary studies examining synaptic morphology, neurotransmitter release, or electrophysiology at defined *C. elegans* synapses would provide additional mechanistic resolution. Finally, while this study characterized four isoform pairs, patients harboring the 3’ deletion express a complex mixture of wild-type and deletion-containing isoforms from the heterozygous locus, and the relative contributions of specific isoforms to neuronal phenotypes in that context remain to be determined. Together, these considerations define a productive roadmap for future work, and the *C. elegans* platform established here provides a genetically tractable foundation for systematic dissection of how individual NRXN1 isoforms contribute to neuronal function and behavioral phenotypes relevant to neuropsychiatric diseases.

## Acknowledgements

The authors thank the labs of John I. Murray, Colin C. Conine, David M. Raizen, and Meera V. Sundaram for their feedback on this project. We also thank members of the Hart lab for comments on the manuscript. Some strains were provided by the CGC, which is funded by NIH Office of Research Infrastructure Programs (P40 OD010440). The authors thank Kristen J. Brennand and Michael B. Fernando for providing plasmids and advice.

## Data and Resource Availability

All relevant data and details of resources can be found within the article and its supplementary information and are available upon request. The data supporting all figures will be posted to Dryad.

## Author Contributions

DH and MPH conceived and designed the study and experiments, and DH conducted all experiments, processed, analyzed, and interpreted all data with help from MPH. DH wrote the manuscript with assistance from MPH.

**Supplementary Table 1.** Strain and plasmid information.

| Strain | Genotype | Co-injection marker |
| --- | --- | --- |
| N2 |  |  |
| MPH232 | *hpmEx38 [pric-19::gfp:: NRXN1 ss39]* | *plin-44::gfp* |
| MPH233 | *hpmEx39 [pric-19::gfp:: NRXN1 ss47]* | *plin-44::gfp* |
| MPH234 | *hpmEx40 [pric-19::gfp:: NRXN1 ss71]* | *plin-44::gfp* |
| MPH235 | *hpmEx41 [pric-19::gfp:: NRXN1 ss75]* | *punc-122::mruby* |
| MPH236 | *hpmEx42 [pric-19::gfp:: NRXN1 ss73]* | *plin-44::gfp* |
| MPH237 | *hpmEx43 [pric-19::gfp:: NRXN1 ss77]* | *plin-44::gfp* |
| MPH238 | *hpmEx44 [pric-19::gfp:: NRXN1 ss84]* | *plin-44::gfp* |
| MPH239 | *hpmEx45 [pric-19::gfp:: NRXN1 ss89]* | *plin-44::gfp* |
| TV13570 | *nrx-1(wy778)* |  |
| MPH240 | *nrx-1(wy778); hpmEx38 [pric-19::gfp:: NRXN1 ss39]* | *plin-44::gfp* |
| MPH241 | *nrx-1(wy778); hpmEx39 [pric-19::gfp:: NRXN1 ss47]* | *plin-44::gfp* |
| MPH242 | *nrx-1(wy778); hpmEx40 [pric-19::gfp:: NRXN1 ss71]* | *plin-44::gfp* |
| MPH243 | *nrx-1(wy778); hpmEx41 [pric-19::gfp:: NRXN1 ss75]* | *punc-122::mruby* |
| MPH244 | *nrx-1(wy778); hpmEx42 [pric-19::gfp:: NRXN1 ss73]* | *plin-44::gfp* |
| MPH245 | *nrx-1(wy778); hpmEx43 [pric-19::gfp:: NRXN1 ss77]* | *plin-44::gfp* |
| MPH246 | *nrx-1(wy778); hpmEx44 [pric-19::gfp:: NRXN1 ss84]* | *plin-44::gfp* |
| MPH247 | *nrx-1(wy778); hpmEx45 [pric-19::gfp:: NRXN1 ss89]* | *plin-44::gfp* |
| MPH49 | *nrx-1(wy778); npr-1(ad609)* |  |
| MPH248 | *nrx-1(wy778); npr-1(ad609); hpmEx38 [pric-19::gfp:: NRXN1 ss39]* | *plin-44::gfp* |
| MPH249 | *nrx-1(wy778); npr-1(ad609); hpmEx39 [pric-19::gfp:: NRXN1 ss47]* | *plin-44::gfp* |
| MPH250 | *nrx-1(wy778); npr-1(ad609); hpmEx40 [pric-19::gfp:: NRXN1 ss71]* | *plin-44::gfp* |
| MPH251 | *nrx-1(wy778); npr-1(ad609); hpmEx41 [pric-19::gfp:: NRXN1 ss75]* | *punc-122::mruby* |
| MPH252 | *nrx-1(wy778); npr-1(ad609); hpmEx42 [pric-19::gfp:: NRXN1 ss73]* | *plin-44::gfp* |
| MPH253 | *nrx-1(wy778); npr-1(ad609); hpmEx43 [pric-19::gfp:: NRXN1 ss77]* | *plin-44::gfp* |
| MPH254 | *nrx-1(wy778); npr-1(ad609); hpmEx44 [pric-19::gfp:: NRXN1 ss84]* | *plin-44::gfp* |
| MPH255 | *nrx-1(wy778); npr-1(ad609); hpmEx45 [pric-19::gfp:: NRXN1 ss89]* | *plin-44::gfp* |
| DA609 | *npr-1(ad609)* |  |

| Plasmid | sequence | Notes |
| --- | --- | --- |
| pMPHDH1 | *pric-19::gfp:: NRXN1 ss39* | * CS-50065 |
| pMPHDH2 | *pric-19::gfp:: NRXN1 ss47* | * CS-50066 |
| pMPHDH3 | *pric-19::gfp:: NRXN1 ss71* | * CS-50067 |
| pMPHDH4 | *pric-19::gfp:: NRXN1 ss75* | * CS-50068 |
| pMPHDH5 | *pric-19::gfp:: NRXN1 ss73* | * CS-50070 |
| pMPHDH6 | *pric-19::gfp:: NRXN1 ss77* | * CS-50074 |
| pMPHDH7 | *pric-19::gfp:: NRXN1 ss84* | * CS-50072 |
| pMPHDH8 | *pric-19::gfp:: NRXN1 ss89* | * CS-50073 |

